# In-bore MRI-compatible Transrectal Ultrasound and Photoacoustic Imaging

**DOI:** 10.1101/2023.11.27.568947

**Authors:** Ryo Murakami, Yang Wang, Wojciech G. Lesniak, Ryosuke Tsumura, Yichuan Tang, Yasuyuki Tsunoi, Christopher J. Nycz, Martin G. Pomper, Gregory S. Fischer, Haichong K. Zhang

## Abstract

Prostate cancer (PCa) is known as one of the most prevalent and fatal cancer types. This report describes an MRI-compatible photoacoustic/ultrasound (PA/US) imaging platform to improve the diagnosis of PCa. In the proposed solution, PA imaging, which offers real-time, non-ionizing imaging with high sensitivity and specificity, is combined with MRI, aiming to overcome PA’s limited field of view (FOV) and make PA scalable for translation to clinical settings. Central to the design of the system is a reflector-based transrectal probing mechanism composed of MRI-compatible materials. The linear transducer with a center hole for optical fiber delivery can be mechanically actuated to form a multi-angled scan, allowing PA/US imaging from varied cross-sectional views. Performance assessment was carried out in phantom and ex-vivo settings. We confirmed the MRI compatibility of the system and demonstrated the feasibility of its tri-modal imaging capability by visualizing a tubing phantom containing contrast agents. The ex-vivo evaluation of targeted tumor imaging capability was performed with a mouse liver sample expressing PSMA-positive tumors, affirming the system’s compatibility in spectroscopic PA (sPA) imaging with biological tissue. These results support the feasibility of the in-bore MRI-compatible transrectal PA and US and the potential clinical adaptability.

## 1. Introduction

Prostate cancer (PCa) continues to be one of the most frequently diagnosed cancer types and ranking as the second leading cause of cancer-related deaths among males [1]. Timely detection of PCa notably increases the effectiveness of treatment and improves patient outcomes. The established diagnostic process for PCa involves screening such as prostate-specific antigen (PSA) tests [2, 3], needle biopsy as definitive diagnosis, and staging evaluation using Gleason score [4]. Imaging techniques including ultrasound (US) and magnetic resonance imaging (MRI) are employed for initial staging, therapy follow-up, and reoccurrence. Positron emission tomography/computed tomography (PET/CT) with prostate-specific membrane antigen (PSMA)-specific radiotracers is a recently introduced alternative [5]. Despite the US offering real-time visualization, it is not tumor or lesion-specific, leading to a systematic biopsy approach, instead of targeted [6]. While MRI and PET/CT are known for their high sensitivity and specificity in PCa imaging [7], they are not typically capable of providing a sufficient frame rate for real-time intraoperative biopsy guidance and often are conducted independently. Given the inherent limitations of current imaging techniques, there is a need for developing an imaging modality that offers non-ionizing, real-time PCa imaging with high sensitivity and specificity.

Photoacoustic (PA) imaging is an emerging non-ionizing imaging technique that integrates the benefits of optical and US imaging [8, 9]. Spectroscopic PA (sPA) provides quantification of multiple indices, which can contribute valuable insights into cancer severity and prognosis. Prostate-specific membrane antigen (PSMA) is a receptor on the surface of PCa cells [10, 11]. It is characterized by its strong correlation with aggressive tumors [12, 13]. In addition to its functional imaging abilities, PA imaging, inclusive of sPA, has been demonstrated to be effective for real-time monitoring [14]. These attributes position PA imaging as a promising modality for PCa diagnosis.

Despite the potential advantages, clinical translatability remains underexplored due to the following limitations. First, the PA imaging system for pre-clinical settings using small animal models [15] is incompatible with humans and large animals, requiring a transrectal configuration. Kothapalli *et al*. presented a transrectal PA imaging apparatus [16]. However, the cross-validation, with other gold-standard imaging modalities is not straightforward because of the tissue deformation and alignment difficulty with data taken in different settings, especially with MRI due to the requirement of the device MRI compatibility. Second, PA imaging has an inconsistent, restricted field of view (FOV) compared to MRI or PET, because the ultrasound receiver geometry determines the imaging field. This makes registering US/PA imaging with other preoperative imaging devices harder. Making the PA-guided procedure to be scalable and compatible with other imaging modalities, especially MRI, could alleviate these constraints.

Here, we propose to develop and MRI-compatible PA/US Imaging Platform. The entire concept is illustrated in Figure 1. In this concept, the PA/US imaging system is placed inside the MRI bore so that these three imaging modalities can be performed without relocating patients. The MRI modality is expected to yield broad FOV images, effectively serving as a global reference for the local imaging supplied by the PA/US imaging. Despite the non-real-time nature of MRI scanning, this approach enables on-demand updates of MRI images, a feature previously unattainable when the PA/US apparatus lacked MRI compatibility, requiring patient movement between the MRI suite and the PA/US room. Leveraging the supplementary role of MRI, it is anticipated that the system can be adapted to larger subjects, facilitating the further exploration of the clinical translatability of PA imaging.

**Figure 1.**
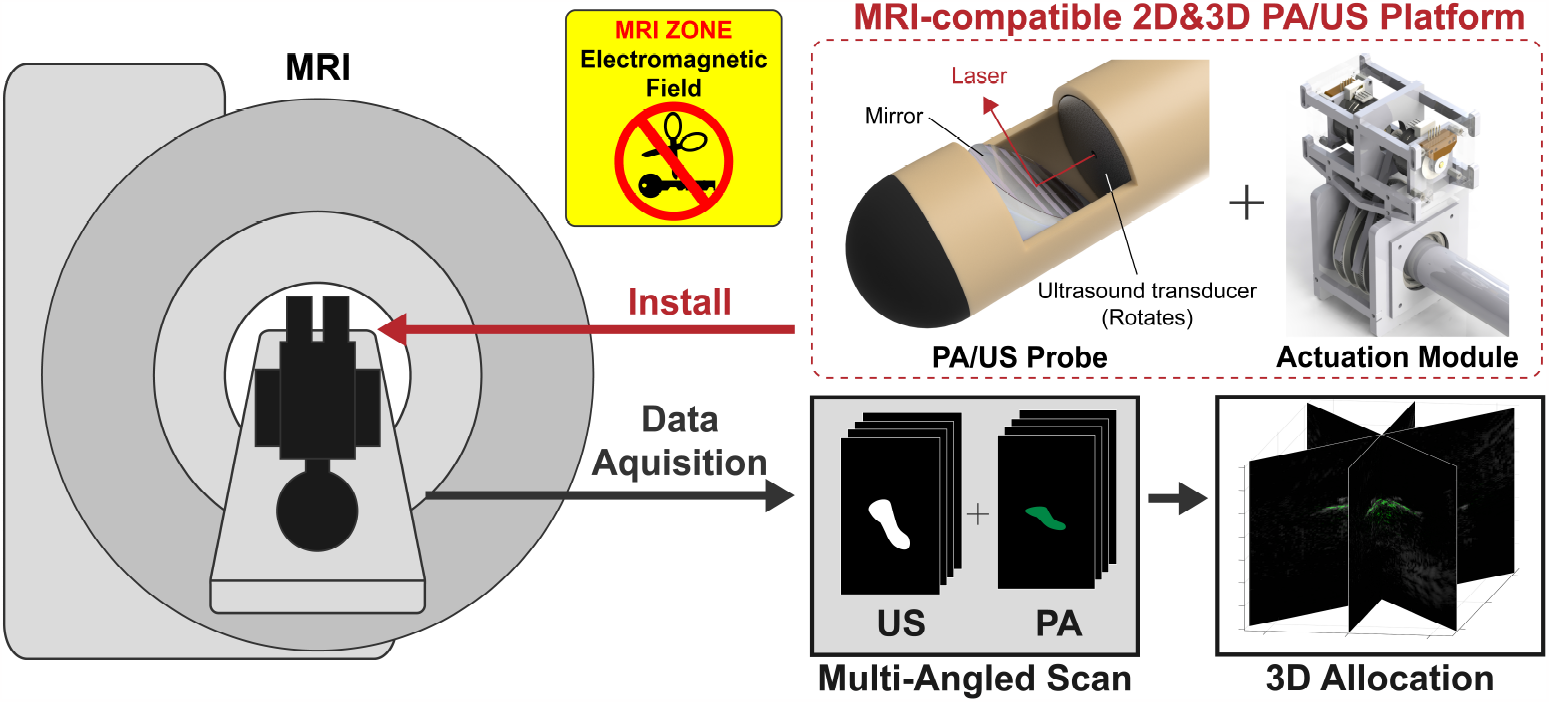
Proposed MRI-compatible PA/US imaging platform and expected operation.

The assurance of MRI compatibility is integral for the realization of our proposed system, as involved in the concept title. Securing this compatibility presents several challenges due to the inherent attributes of the MRI scanner, including the magnetic field, transitioning magnetic field gradients, and radio frequency pulses, as well as the MRI’s sensitivity to external interference [17]. For example, the strict prohibition of magnetic materials and the requirement to reduce non-magnetic metallic components to avoid potential risks like induced thermal effects and artifacts in MRI images. The domain of MRI-compatible devices, such as robotics, has witnessed considerable research efforts, with several groups focusing on addressing these limitations [18, 19, 20, 21, 22]. In the specific context of MRI-compatible PA device development, Chen, Gezginer *et al*. developed an MRI-compatible PA tomography system for pre-clinical imaging studies [23, 24].

In this report, we introduce a design for an MRI-compatible PA/US imaging platform. This platform integrates a custom-made MRI-compatible PA/US probe with its associated actuation module. The performance of the newly developed system is evaluated using a phantom model and an *ex-vivo* setup.

The subsequent portions of this manuscript are structured in the following manner: Section 2 primarily covers the design of the MRI-compatible PA/US probe, its actuation module, along with the experimental frameworks, and the parameters for the following evaluations. The observed findings are compiled and explained in Section 3. Section 4 discusses the performance of the established system, potential applications, and constraints rooted based on the obtained results. Lastly, Section 5 concludes the entire paper.

## 2. Materials and Methods

### 2.1. Customized MRI-Compatible PA/US Probe

A customized MRI-compatible PA probe is developed (Figure 2). It consists of two parts: the outer shaft and the inner shaft. The outer shaft is made of ULTEM1010 (ULTEM 1010, Natural, Solid Infill, Fused Deposition Modeling (FDM), Xeometry, Maryland, United States), and the inner shaft is made of brass. Between the two shafts, bearings made of glass-filled PTFE (iglide^®^ i3-PL, igus, Inc., Rhode Island, United States) are placed to smoothen the rotation. On the distal side of the inner shaft, a 68-element linear array transducer (68-element linear array transducer (element pitch: 0.2 mm), Japan Probe Co, Ltd., Kanagawa, Japan) is fixed and rotates with the inner shaft. The transducer has a hole (diameter: 2 mm) at the center of the array so that an optical fiber (FT1000EMT, Thorlabs, New Jersey, United States) can pass through it and shoot a laser. A dielectric mirror (25 mm Diameter 750 - 1100 nm Broadband *λ*/10 Mirror, Edmund Optics Inc., New Jersey, United States) stand is fixed to the outer shaft and the mirror has a 45-degree slope with respect to the horizontal axis. The mirror is designed so that it can reflect both the acoustic wave and the laser, altering their trajectories by 90 degrees.

**Figure 2.**
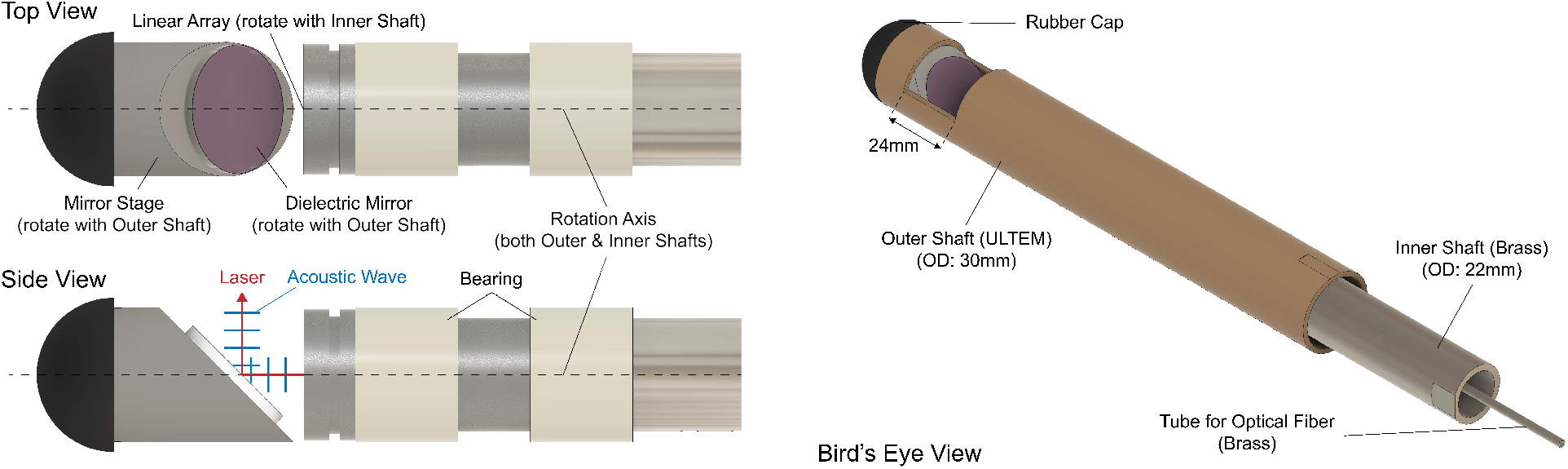
Developed MRI-compatible PA/US probe

### 2.2. Probe Actuation System

An MRI-compatible probe actuation system is developed and fabricated with MRI-safe material and MRI-conditional electronics. Figure 3 represents the CAD model of the developed actuation module. A dedicated control box and communication flow are also designed for the module.

**Figure 3.**
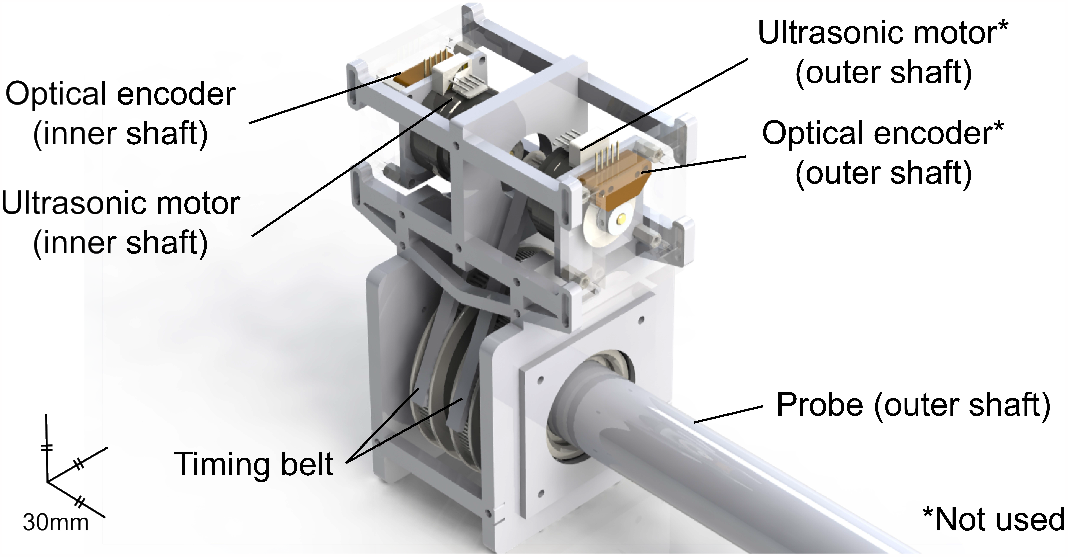
CAD model of the actuation module with PA Probe Mounted

An ultrasonic motor (USR30-S4N, SHINSEI Corporation, Tokyo, Japan) is used as an actuator, ensuring its compatibility with MRI by precluding the use of magnetic materials. A timing belt system is used to transfer the movement from the motor to the inner shaft, with a predetermined ratio of 5: 1. A 1250 CPR optical encoder (E2-1250-157-IE-H-D-1, US Digital Corporation, WA, United States) is mounted on the back shaft of the motor to monitor the rotation of the shaft, which is converted corresponding to the predetermined ratio.

### 2.3. Multi-Angled Scanning

The developed imaging probe is designed so that the rotation of the linear transducer enables the multi-angled scan of 2D PA/US images (Figure 4 (a)). The acquired imaging slices are allocated to their corresponding angles (Figure 4 (b)). Here, *i* is the transducer element number from the center of rotation and *j* is the slice number. Since the linear array has 68 elements in total, the maximum number of *i* is 34. The three-dimensional data is generated in the MATLAB environment (MATLAB, The MathWorks, Inc., Massachusetts, United States). As the number of imaging slices for this scanning increases, the distance between the slices becomes smaller, determining the scanning density in the 3D space. During the rotation, the mirror attached to the outer probe shaft is fixed. In PA, 64 frames are obtained for each slice and wavelength, and the total number of wavelengths applied is six (730, 750, 780, 800, 820, and 850 nm).

**Figure 4.**
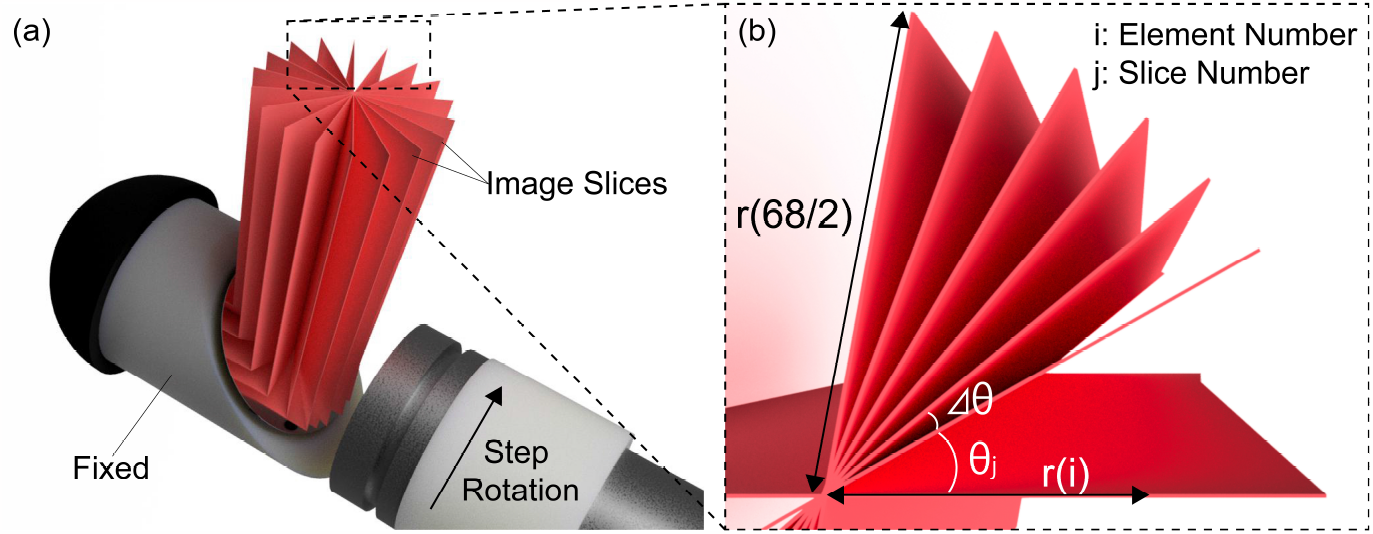
Schematics of the targeted 3D image scanning & reconstruction. (a) Bird’s-eye view, (b) Top view of the image slices.

### 2.4. Spectroscopic Decomposition

For the purpose of functional PA imaging, a technique called spectroscopic decomposition is utilized in this study. The received PA signals can be assumed as linear combinations of multiple absorbers in a tissue or contrast agents; therefore, the obtained signals can be decoupled into the contribution of each absorber by referring to their absorbing characteristics. This concept can be formularized as shown in Equation 1.

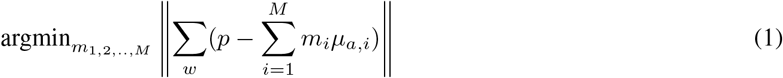

where *p* is the obtained PA spectrum, *μ*_*a, i*_ and *M* are the absorption spectrum of the contrast *i* and *M* is the number of assumed optical absorbers. *w* is the applied laser wavelength. *m*_*i*_ as the weight of the contract *i*, is calculated as an output of this equation [13].

### 2.5. Experimental Setup

#### 2.5.1. System Architecture

Figure 5 shows the system architecture. The PA/US probe, a target sample, and the actuation module consisting of the ultrasonic motors and the encoders are placed in the MRI bore (SIGNA Premier 3.0T MRI scanner, GE Healthcare Technologies Inc., Chicago, IL, United States), and the control box includes a power source, drivers, a controller board. As the laser device (Phocus MOBILE, OPOTEK, Inc., Carlsbad, CA, United States) is used and the ultrasound data acquisition system (Vantage 128, Verasonics, Kirkland, WA, United States) is located in the closet room, which is next to the MRI room and shielded from the MRI-induced electromagnetic field. The cables required for the actuation module and transducer, along with the optical fiber necessary for the laser system, are passed through the brass partition that separates the equipment room from the MRI room.

**Figure 5.**
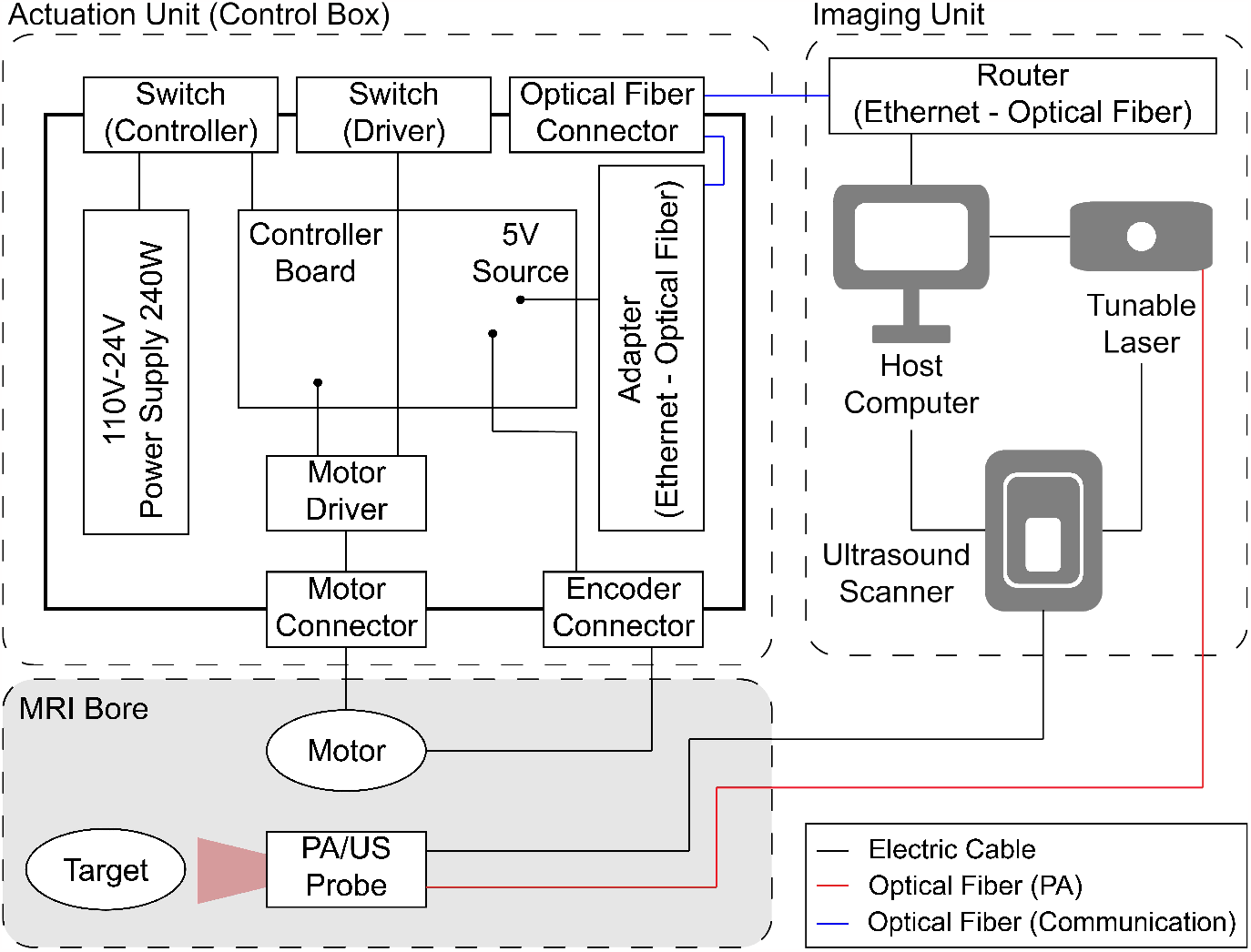
System Architecture involving Actuation Unit and Imaging Unit.

A control box is comprised of the controller (DMC-4143-CARD, Galil Motion Control, Inc., Rocklin, CA, United States) for the actuation module, the drivers (D6030/24V, SHINSEI Corporation, Tokyo, Japan) for the motors, the media converter that transforms optical signal to serial signal, and two power supplies. The communication between the control box and the computer is established using a group of optical fiber cables, a media converter, and a router that acts as the hub of the communication network. The control box was positioned in the equipment room, rather than the MRI room for safety in this particular experiment; however, the optical communication system is designed with the intent of allowing the control box to be located in the MRI room, thereby potentially facilitating additional noise reduction.

#### 2.5.2. Sample Preparation

- Wire Phantom (3 × 2 grid) A schematic representation of the experimental setup for a signal-to-noise ratio (SNR) and resolution analysis with a wire phantom is presented in Figure 6 (a). The medium employed in this setup is water. The wire phantom consists of six parallel-aligned black wires (Diameter: 0.2 mm), which serve as point targets. In the horizontal and vertical directions, the distance between adjacent wires is set at 5 mm and 10 mm, respectively.

**Figure 6.**
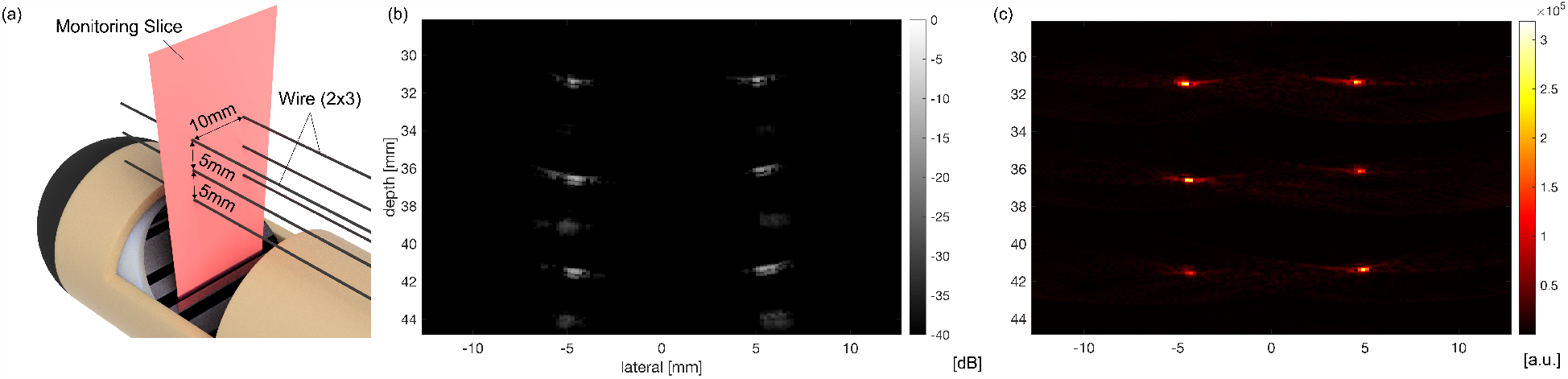
(a) Experimental setup of the wire phantom, (b) US image, (c) Single-wavelength (710 nm) PA image.
- Tube Phantom with Contrast Agents Three tubing targets are prepared to evaluate the MRI compatibility of the developed system and its tri-modal imaging capability. It employs indocyanine green (ICG) solution (Indocyanine green 250 mg, U.S. Pharmacopeia, United States) and MR-SPOT^®^, radiation water (MR-SPOT^®^ 121, Beekley Medical^®^, Connecticut, United States). Each silicone tube (Outer diameter (OD): 2 mm, Inner diameter (ID): 1 mm) encapsulates either 25*μ*M ICG solution or MR-SPOT^®^ exclusively, or a 2 : 1 mixture of 25*μ*M ICG and MR-SPOT^®^. The configuration of the tubes is depicted in Figure 7 (a) and their cross-sections are scanned with MRI, US, and PA employing ultrasound gel as the coupling medium. With this setup, it can be hypothesized that solely PA imaging can distinguish the tubes containing ICG based on the ICG-specific optical absorbance characteristics. The wavelengths shot for this phantom range from 700 to 850 nm with the 10 nm steps.

**Figure 7.**
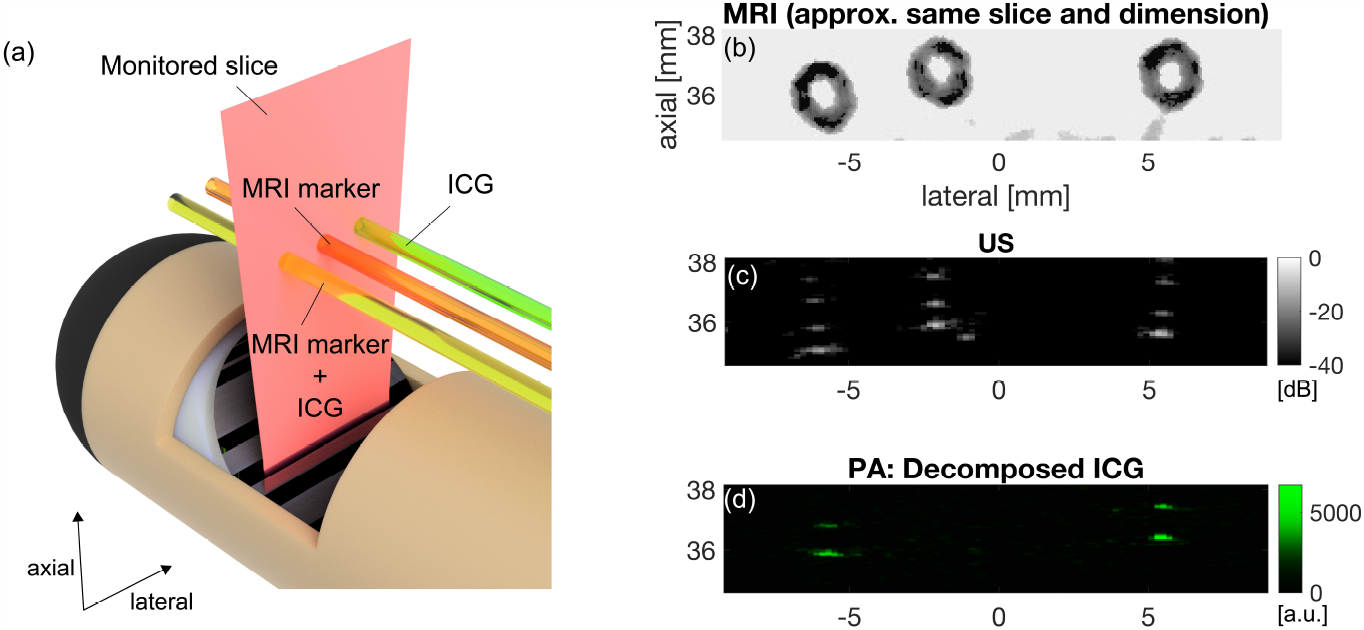
(left, a) Configuration of the tube phantom consisting of 25 *μ*M ICG, MRI marker, Mixture of 25 *μ*M ICG + MRI marker (mixing ratio: 2 : 1), (right) Result of the two-dimensional PA/US imaging and the MRI scan. ((b) MRI image, (c) US, (d) Decomposed ICG in sPA.
- *ex-vivo* Sample (Mouse liver with PCa cells containing contrast agent) An *ex-vivo* sample consisting of a mouse liver containing a contrast agent, Conjugate IV described earlier [25], is utilized (Figure 9 (a)). The agent was designed for the visualization of PSMA in PCa with PA and fluorescence. The imaging agent was synthesized trough consecutive conjugation on average 6 sulfo-Cy7.5 near infrared dyes, 9 suberic acid-lysine-glutamate-urea (SA-KEU) PSMA targeting moieties and 53 butane-1, 2-diol capping moieties with generation-4 amine terminated poly (amidoamine) dendrimer (G4(Cy7.5)_6_(SA-KEU)_9_(Bdiol)_53_ [25]. The sample was prepared and provided by the group of Johns Hopkins University and preserved at around −80^*o*^C at a designated freezer and subsequently transferred to another freezer with a higher temperature one day prior to the imaging session. During the experiment procedure, the sample’s position and orientation were stabilized using ultrasound gel, which served as the coupling medium.

### 2.6. Evaluation Metrics

#### 2.6.1. Indexes for Image Quality Evaluation

To quantify the quality of the acquired two-dimensional images, a SNR and a resolution are computed in this study by visualizing the grid wire phantom. Resolution is computed as the full width at half maximum (FWHM). SNR is defined as shown in the following Equation 2.

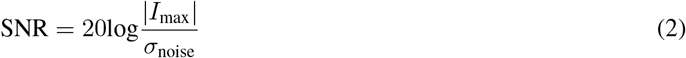

where *I*_max_ is the peak signal intensity at a target and *σ*_noise_ represents the standard deviation of signal intensities in an experimentally defined noise area [26].

#### 2.6.2. Evaluation Criteria for MRI Compatibility

In accordance with the criteria for MRI compatibility as delineated by Tsekos *et at*. [17], the criteria for a device to be deemed “MRI compatible” within the context of this research are defined as follows with the designed experiment for each evaluation:

Criterion 1: The device can be safely placed in the MRI environment. (e.g., magnetic force)

- The probe and the actuation module are placed inside the MRI bore and see confirm if any force is induced.

Criterion 2: The presence of the device does not significantly affect the MRI image quality.

- The MRI imaging is conducted around the image slice of PA/US with the probe and the actuation module placed inside the MRI bore.

Criterion 3: The operation of the device is not affected by the MRI scanner.

- PA/US imaging and actuation are performed inside the MRI and their performance is evaluated.

## 3. Results

### 3.1. Resolution and SNR Evaluation

We evaluated the image quality of US and PA imaging using the wire phantom (Figures 6 (b) and (c)) as the imaging performance of the developed system. In both imaging methods, six-point targets as the cross-section of wires were visualized. Table 1 summarizes the quantified FWHM and SNR. The indexes are evaluated for each point target, then the mean and the standard deviation values are computed for each depth (Table 1).

**Table 1:** Lateral resolution (FWHM) and SNR for each depth (approx.) based on wire phantom.

| Depth [mm] | FWHM [mm] |  |  | SNR [dB] |  |  |
| --- | --- | --- | --- | --- | --- | --- |
|  | 31 | 36 | 41 | 31 | 36 | 41 |
| US | $1.30 \pm 0.14$ | $2.10 \pm 0.42$ | $3.50 \pm 0.71$ | $48.8 \pm 6.72$ | $48.98 \pm 5.26$ | $52.45 \pm 3.99$ |
| PA | $1.80 \pm 0.00$ | $1.80 \pm 0.00$ | $2.10 \pm 0.14$ | $39.17 \pm 1.94$ | $37.91 \pm 3.38$ | $40.40 \pm 1.18$ |

### 3.2. MRI Compatibility Evaluation

In order to confirm the MRI compatibility of the developed system, including the imaging probe and the actuation module, the evaluation was performed along with the three criteria as defined in Section 2.6.2.

#### Criterion 1: Safe installation

It is confirmed that the safe installation of the developed system, including the PA/US probe and the actuation module, into the MRI bore, and no significant movements of the system attributable to the MRI scanner were observed throughout the experiment. Consequently, we can affirm that the first criterion for MRI compatibility has been successfully fulfilled.

#### Criterion 2: MRI image quality

To evaluate how much the developed system affects the quality of MRI images, the MRI image was obtained. Artifacts caused by the metals involved in the developed system were observed, these were sufficiently minimal to enable visualization of targets at around 10 mm or greater from the outer shaft wall of the probe. This result suggests that Criterion 2 is met, given that a target maintains a specific distance from the probe’s outer shaft wall to avoid artifact interference.

#### Criterion 3: System performance

For this criterion, both the performance of the actuation module and the imaging unit need to be evaluated. The actuation module was controlled inside the MRI bore to evaluate its performance in the MRI environment. As a control target, 1 deg/step up to 180 degrees was applied to the inner shaft. In the comparison between the target angles and the measured angles with the encoder (Figure 8), the error in each rotation step was 0.03 *±* 0.01 deg.

**Figure 8.**
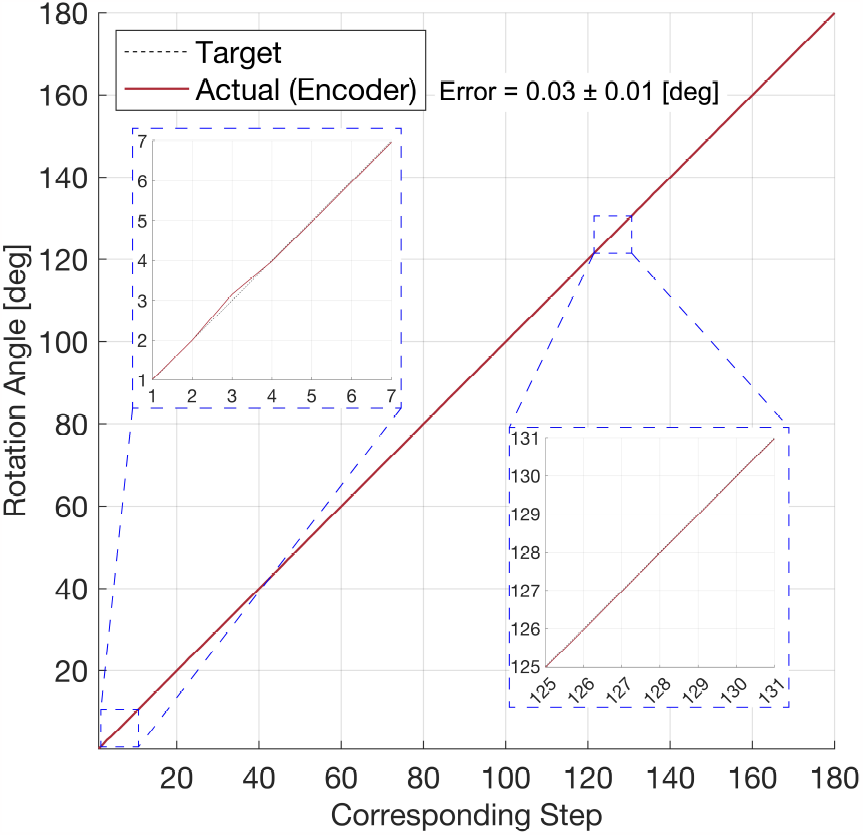
Result of actuation accuracy evaluation for MRI compatibility.

**Figure 9.**
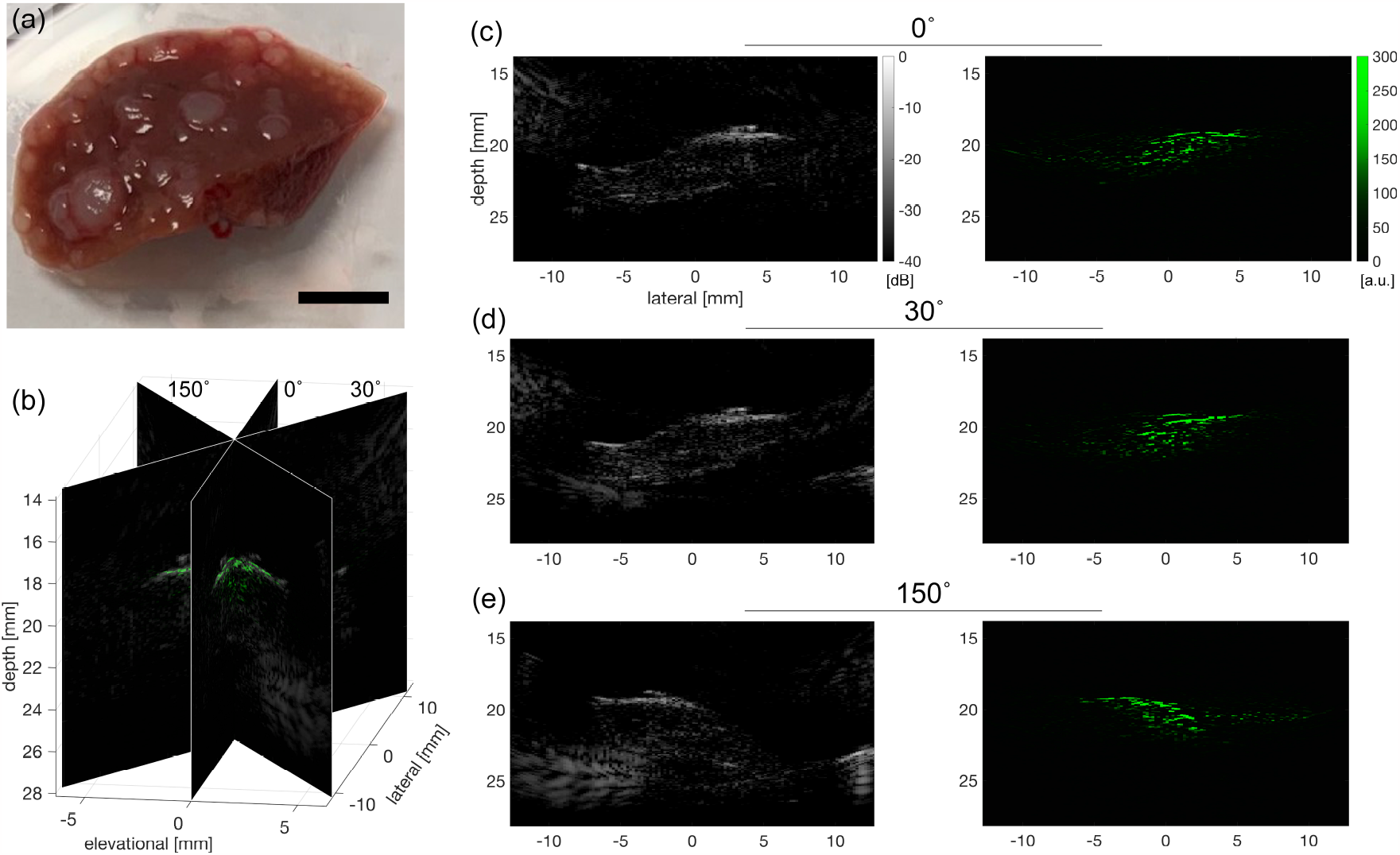
Result of *ex-vivo* imaging capability evaluation; (a) Photograph of the liver sample containing PCa cells and contrast agent (Scale: 5 mm), (b) Multi-angled imaging slices allocated to their corresponding angles, (c-e) 2D US and contrast agent-targeted sPA for each angle (0^*o*^, 30^*o*^, and 150^*o*^).

Also, considering the encoder resolution and the transmission ratio, the measurement resolution with the encoder is computed as 0.009 deg in this configuration. Regarding the imaging performance, US and PA imaging results are demonstrated in Figure 7 (c) and (d), respectively, verifying that each imaging modality yields plausible outcomes. These results support the fact that both the PA/US imaging and the actuation do work under the MRI environment, suggesting the system satisfies Criterion 3.

Since the three criteria for MRI compatibility have been satisfied as discussed, it can be concluded that the MRI compatibility of the developed system with the configuration is confirmed based on the definition.

### 3.3. Tri-Modal Imaging Capability Evaluation

For the evaluation of the tri-modal imaging capability of the entire system, PA, US, and MRI imaging were performed (Figure 7 (b-d)). The PA/US probe and the actuation module were placed inside the MRI bore throughout all the scans. The registration of the MRI image to PA/US was performed based on the geometrical relationship between the probe geometry captured by MRI and the PA/US imaging window. Once the MRI slice is found, the image is manually scaled to be aligned with the US image. Although both MRI and US can depict all the tube cross-sections, the circles with holes are most clearly revealed in the MRI image. The US image captures the inner and outer surface of tubings via acoustic reflection, while the PA image depicts signals from the contrast agent appearing at the inner surface. Taking into account the geometry of the tube (OD: 2 mm, ID: 1 mm), it is confirmed that the registered scale of the MRI image is appropriate by comparing it with the scales of PA and US.

As demonstrated in Figure 7, MRI, US, and PA imaging were achieved without any change in the pose of the imaging target and the imaging device. This eliminates potential organ deformation and alignment complications, realizing more reliable image registration. Moreover, the MRI image’s wide FOV in three dimensions facilitates the detection of the probe’s geometry, enabling geometry-based registration. Hence, the proposed MRI-compatible PA/US imaging platform addressed the limitations of PA imaging alone mentioned in Section 1.

Spectroscopic decomposition with respect to the ICG spectrum was carried out to assess the performance of sPA imaging, as shown in Figure 7 (d). The sPA result distinctly emphasizes the two tubes holding the ICG solution. This result demonstrates the device’s ability for selective visualization of PCa, which is the expected function of PA imaging for this application.

### 3.4. *ex-vivo* Sample Imaging Capability Evaluation with Multi-Angled Scanning

The objectives of this ex-vivo study comprise evaluating the PA/US imaging capability on biological tissue and the feasibility of the multi-angled scanning capability, which is the simplified version of the 3D scanning with 1 deg/step angle resolution. The two-dimensional PA/US imaging outcomes of the *ex-vivo* tumor sample containing Conjugate IV as a contrast agent are depicted (Figure 9 (c: 0^*o*^), (d: 30^*o*^), and (e: 150^*o*^)). The anatomical feature of the sample is delineated in the US image, and the Conjugate IV-targeted sPA returns the corresponding signal from the sample. Each 2D image has the corresponding angle of the transducer as metadata. Therefore, these planes can be displayed in three-dimensional space based on their angles (Figure 9 (b)). Considering the PSMA-targeted PA contrast agent attaches tumors in the liver sample comprising human PCa cells, this *ex-vivo* imaging evaluation appears suitable for the PA/US imaging device designed specifically for PCa imaging.

## 4. Discussion

This research investigated the MRI-compatible transrectal US and PA imaging system, including its actuation module, considering its application to PCa. The performance of this system has been evaluated within both the phantom and the ex-vivo settings.

The PA/US probe engineered utilizes a reflector-based mechanism to facilitate transrectal PA/US, an approach corroborated in the US imaging by Tsumura *et al*. [26]. Their research verified that the reflector’s presence does not substantially degrade image quality, as evidenced by a comparative analysis of images taken with and without the reflector. In the current study, the PA functionality was introduced through the integration of an optical fiber.

Given that the incorporated dielectric mirror consistently maintains an average reflection rate exceeding 96% within the wavelength range used in this experiment, it suggests that the PA imaging quality is not significantly compromised by the reflector.

During the phantom experiment, tubings were positioned as such to avoid artifact interference in the MRI image from the developed hardware, leading to its successful visualization in the MRI image. The majority of the artifact appears to be associated with the brass material around the transducer. The replacement of the brass material with a non-metallic substrate, such as ULTEM used in the outer shaft, is anticipated to curtail the artifact. As the underlying concept will be implemented in real-world clinical contexts, minimizing such artifacts and securing an expansive FOV becomes essential.

One of the core aims of the present study is to facilitate accompanying target visualization of PA, US, and MRI, with no requirement for patient movement between scans. Besides the necessary MRI compatibility of the PA/US probe, a fundamental prerequisite for the system is its capacity for image “FUSION” - the registration of all three imaging modalities in terms of spatial position and orientation. Several potential methodologies exist for image registration, one of which involves identifying the PA/US imaging slice within the MRI image, informed by the geometric characteristics of the probe as visualized via MRI. The addition of reference points, such as an MRI marker, may aid in discerning the geometric relationship between the probe and the MRI image. In this particular investigation, the first approach was used, and the MRI image displayed in Figure 7 (b) was manually scaled and aligned by referencing the corresponding US image (Figure 7 (c)). This process could be automated, as demonstrated in some existing MRI-US FUSION platforms [27], provided the relevant MRI imaging plane is identified.

While the proposed system exhibited anticipated performance characteristics, several limitations should be considered. First, water or transparent ultrasound gel was utilized as the acoustic coupling medium throughout the experiments. This circumstance offers a more advantageous scenario than that provided by actual biological tissues in terms of depth penetration, especially for PA. Towards the translation into real-world clinical settings, improving the PA’s depth penetration is crucial to secure a wider FOV. Potential countermeasures include adding an optical fiber independently from the transducer, passing it through the urethra for instance, and directing a laser proximal to the target, or increasing the core diameter of the optical fiber to facilitate the delivery of enhanced laser energy. Second, the 3D scanning capability of both PA and US needs a more thorough investigation, although the feasibility of the multi-angled scan has been confirmed. To evaluate and validate the 3D scanning capability, the 3D scan with 1 deg/step resolution for a geometry-known phantom should be performed as one of the future works.

## 5. Conclusions

This research introduces the concept of an MRi-compatible PA/US imaging platform with the aim of tri-modal imaging, enhancing the scalability of PA imaging and encouraging further examination of its clinical adaptability by incorporating MRI imaging as a supplementary modality. An MRI-compatible transrectal PA/US probe was developed, together with its dedicated actuation system. The MRI compatibility and the feasibility of the proposed tri-modal imaging have been evaluated under the phantom environment. This demonstrates its safe and effective operation within the MRI environment, including sPA imaging. The *ex-vivo* study underscored the system’s ability to capture images of biological tissue in both multi-angled imaging formats. Future research will focus on improving the system’s reliability and stability, as well as conducting *in-vivo* studies.

## Acknowledgements

This work was supported by National Institutes of Health under grants CA134675, DK133717, and OD028162.

## Declaration

During the preparation of this work, the authors used ChatGPT for proofreading in order to enhance the readability and eliminate grammatical errors. After using this tool, the authors reviewed and edited the content and take full responsibility for the content of the publication.

## References

[1] L. Siegel Rebecca, D. Miller Kimberly, A. Jemal, Cancer statistics, 2016, CA Cancer J Clin 66 (1) (2016) 7–30.

[2] D. Ilic, M. Djulbegovic, J. H. Jung, E. C. Hwang, Q. Zhou, A. Cleves, T. Agoritsas, P. Dahm, Prostate cancer screening with prostate-specific antigen (psa) test: a systematic review and meta-analysis, bmj 362 (2018).

[3] J. H. Hayes, M. J. Barry, Screening for prostate cancer with the prostate-specific antigen test: a review of current evidence, Jama 311 (11) (2014) 1143–1149.

[4] A. W. Partin, J. Yoo, H. B. Carter, J. D. Pearson, D. W. Chan, J. I. Epstein, P. C. Walsh, The use of prostate specific antigen, clinical stage and gleason score to predict pathological stage in men with localized prostate cancer, The Journal of urology 150 (1) (1993) 110–114.

[5] E. M. Schaeffer, S. Srinivas, N. Adra, Y. An, D. Barocas, R. Bitting, A. Bryce, B. Chapin, H. H. Cheng, A. V. D’Amico, et al., Nccn guidelines® insights: Prostate cancer, version 1.2023: Featured updates to the nccn guidelines, Journal of the National Comprehensive Cancer Network 20 (12) (2022) 1288–1298.

[6] D. Piao, Z. Jiang, K. E. Bartels, G. R. Holyoak, J. W. Ritchey, K. Rock, C. L. Ownby, C. F. Bunting, G. Slobodov, Optical biopsy of the prostate: can we TRUST (trans-rectal ultrasound-coupled spectral tomography)?, in: R. R. Alfano (Ed.), Optical Biopsy IX, Vol. 7895, International Society for Optics and Photonics, SPIE, p. 78950I.

[7] K.-H. Pan, J.-F. Wang, C.-Y. Wang, A. A. Nikzad, F. Q. Kong, L. Jian, Y.-Q. Zhang, X.-M. Lu, B. Xu, Y.-L. Wang, et al., Evaluation of 18f-dcfpyl psma pet/ct for prostate cancer: a meta-analysis, Frontiers in Oncology 10 (2021) 597422.

[8] M. Xu, L. V. Wang, Photoacoustic imaging in biomedicine, Review of Scientific Instruments 77 (4) (2006) 041101. doi:10.1063/1.2195024.

[9] L. V. Wang, Prospects of photoacoustic tomography, Medical Physics 35 (12) 5758–5767.

[10] X. Huang, M. Bennett, P. E. Thorpe, Anti-tumor effects and lack of side effects in mice of an immunotoxin directed against human and mouse prostate-specific membrane antigen, The Prostate 61 (1) (2004) 1–11.

[11] N. Schülke, O. A. Varlamova, G. P. Donovan, D. Ma, J. P. Gardner, D. M. Morrissey, R. R. Arrigale, C. Zhan, A. J. Chodera, K. G. Surowitz, et al., The homodimer of prostate-specific membrane antigen is a functional target for cancer therapy, Proceedings of the National Academy of Sciences 100 (22) (2003) 12590–12595.

[12] S. Perner, M. D. Hofer, R. Kim, R. B. Shah, H. Li, P. Möller, R. E. Hautmann, J. E. Gschwend, R. Kuefer, M. A. Rubin, Prostate-specific membrane antigen expression as a predictor of prostate cancer progression, Human pathology 38 (5) (2007) 696–701.

[13] H. K. Zhang, Y. Chen, J. Kang, A. Lisok, I. Minn, M. G. Pomper, E. M. Boctor, Prostate-specific membrane antigen-targeted photoacoustic imaging of prostate cancer in vivo, Journal of Biophotonics 11 (9) e201800021. doi:10.1002/jbio.201800021.

[14] R. Murakami, S. Gao, H. K. Zhang, Closed-loop contrast source regulation through real-time spectroscopic photoacoustic imaging: phantom evaluation, in: Photons Plus Ultrasound: Imaging and Sensing 2023, Vol. 12379, SPIE, 2023, pp. 265–270.

[15] N. Singh, E. Cherin, Y. Soenjaya, A. Patel, C.-F. Roa, G. Zheng, F. S. Foster, C. E. Demore, Photoacoustic imaging of a pre-clinical tumour model with an exactvu micro-ultrasound system, in: 2022 IEEE International Ultrasonics Symposium (IUS), IEEE, 2022, pp. 1–4.

[16] S.-R. Kothapalli, G. A. Sonn, J. W. Choe, A. Nikoozadeh, A. Bhuyan, K. K. Park, P. Cristman, R. Fan, A. Moini, B. C. Lee, et al., Simultaneous transrectal ultrasound and photoacoustic human prostate imaging, Science translational medicine 11 (507) (2019) eaav2169.

[17] N. V. Tsekos, A. Khanicheh, E. Christoforou, C. Mavroidis, Magnetic resonance–compatible robotic and mechatronics systems for image-guided interventions and rehabilitation: a review study, Annu. Rev. Biomed. Eng. 9 (2007) 351–387.

[18] G. S. Fischer, I. Iordachita, C. Csoma, J. Tokuda, S. P. DiMaio, C. M. Tempany, N. Hata, G. Fichtinger, Mri-compatible pneumatic robot for transperineal prostate needle placement, IEEE/ASME transactions on mechatronics 13 (3) (2008) 295–305.

[19] A. Krieger, S.-E. Song, N. B. Cho, I. I. Iordachita, P. Guion, G. Fichtinger, L. L. Whitcomb, Development and evaluation of an actuated mri-compatible robotic system for mri-guided prostate intervention, IEEE/ASME Transactions on Mechatronics 18 (1) (2011) 273–284.

[20] K. Masamune, E. Kobayashi, Y. Masutani, M. Suzuki, T. Dohi, H. Iseki, K. Takakura, Development of an mri-compatible needle insertion manipulator for stereotactic neurosurgery, Journal of image guided surgery 1 (4) (1995) 242–248.

[21] K. Chinzei, R. Kikinis, F. A. Jolesz, Mr compatibility of mechatronic devices: design criteria, in: Medical Image Computing and Computer-Assisted Intervention–MICCAI’99: Second International Conference, Cambridge, UK, September 19-22, 1999. Proceedings 2, Springer, 1999, pp. 1020–1030.

[22] J. F. de Groot, A. H. Kim, S. Prabhu, G. Rao, A. W. Laxton, P. E. Fecci, B. J. O’Brien, A. Sloan, V. Chiang, S. B. Tatter, A. M. Mohammadi, D. G. Placantonakis, R. E. Strowd, C. Chen, C. Hadjipanayis, M. Khasraw, D. Sun, D. Piccioni, K. D. Sinicrope, J. L. Campian, S. C. Kurz, B. Williams, K. Smith, Z. Tovar-Spinoza, E. C. Leuthardt, Efficacy of laser interstitial thermal therapy (LITT) for newly diagnosed and recurrent IDH wild-type glioblastoma, Neuro-Oncology Advances 4 (1) (04 2022). doi:10.1093/noajnl/vdac040.

[23] Z. Chen, I. Gezginer, M.-A. Augath, W. Ren, Y.-H. Liu, R. Ni, X. L. Deán-Ben, D. Razansky, Hybrid magnetic resonance and optoacoustic tomography (mrot) for preclinical neuroimaging, Light: Science & Applications 11 (1) (2022) 332.

[24] I. Gezginer, Z. Chen, H. Yoshihara, X. L. Deán-Ben, D. Razansky, Volumetric registration framework for multimodal functional magnetic resonance and optoacoustic tomography of the rodent brain, Photoacoustics (2023) 100522.

[25] W. G. Lesniak, Y. Wu, J. Kang, S. Boinapally, S. R. Banerjee, A. Lisok, A. Jablonska, E. M. Boctor, M. G. Pomper, Dual contrast agents for fluorescence and photoacoustic imaging: evaluation in a murine model of prostate cancer, Nanoscale 13 (20) (2021) 9217–9228.

[26] R. Tsumura, Y. Tang, H. K. Zhang, Reflector-based transrectal 3d ultrasound imaging system for transperineal needle intervention, in: 2020 IEEE International Ultrasonics Symposium (IUS), 2020, pp. 1–4. doi:10.1109/IUS46767.2020.9251655.

[27] J. K. Logan, S. Rais-Bahrami, B. Turkbey, A. Gomella, H. Amalou, P. L. Choyke, B. J. Wood, P. A. Pinto, Current status of magnetic resonance imaging (mri) and ultrasonography fusion software platforms for guidance of prostate biopsies, BJU international 114 (5) (2014) 641–652.

